# Upregulation of TRPM3 drives hyperexcitability in nociceptors innervating inflamed tissue

**DOI:** 10.1101/2020.04.30.069849

**Authors:** Marie Mulier, Nele Van Ranst, Nikky Corthout, Sebastian Munck, Pieter Vanden Berghe, Joris Vriens, Thomas Voets, Lauri J Moilanen

## Abstract

Genetic ablation or pharmacological inhibition of the heat-activated cation channel TRPM3 alleviates heat hyperhyperalgesia in animal models of inflammation, but the mechanisms whereby the channel contributes to inflammatory pain are unknown. Here, we induced unilateral inflammation of the hind paw in mice, and directly compared expression and function of TRPM3 and two other heat-activated TRP channels (TRPV1 and TRPA1) in sensory neurons innervating the ipsilateral and contralateral paw. We detected increased Trpm3 mRNA levels in dorsal root ganglion neurons innervating the inflamed paw, as well as augmented TRP channel-mediated calcium responses, both in the cell bodies and the intact peripheral endings of nociceptors. Notably, inflammation provoked a pronounced increase in nociceptors co-expressing functional TRPM3 with TRPV1 and TRPA1, and pharmacological inhibition of TRPM3 caused normalization of TRPV1- and TRPA1-mediated responses. These new insights into the mechanisms underlying inflammatory heat hypersensitivity provide a rationale for developing TRPM3 antagonists to treat pathological pain.

## Introduction

Painful stimuli are detected by nociceptors in peripheral tissues and are transmitted as action potentials towards the central nervous system to elicit pain sensation (1–3). Under pathological conditions, such as inflammation or tissue injury, nociceptors become sensitized to mechanical and thermal stimuli. Such nociceptor hypersensitivity gives rise to allodynia (the sensation of pain to a stimulus that is usually not painful), hyperalgesia (increased pain sensation to a stimulus that is usually painful) or to spontaneous pain without any clear stimulus (1, 2, 4). The current analgesic drug therapies, including non-steroidal anti-inflammatory drugs, opioids, gabapentinoids and antidepressants, are often limited in efficacy in a large number of pain patients, and may have severe adverse effects that limit their use (5–7). The need for better and safer analgesic drugs is painfully illustrated by the dramatic rise in opioid addiction and related deaths, known as the opioid crisis (6). The search for new analgesic drugs with novel mechanisms of action inevitably depends on a deep understanding of the cellular and molecular mechanisms underlying nociceptor sensitization (5).

In this context, several members of the transient receptor potential (TRP) superfamily of cation channels play key roles as primary molecular sensors in nociceptor neurons, directly involved in translating external stimuli into neuronal activity and pain (8). For instance, we recently demonstrated that heat-induced pain in mice depends on a trio of heat-activated TRP channels, TRPM3, TRPA1 and TRPV1 (9). Robust neuronal and behavioral heat responses were observed as long as at least one of these three TRP channels was functional, but were fully abolished in *Trpm3*^*−/−*^/*Trpv1*^*−/−*^/*Trpa1*^*−/−*^ triple knockout mice (9). These results indicate triple redundancy of the molecular sensors for acute heat. Intriguingly, however, there apparently is no such redundancy for the development of heat hypersensitivity in inflammatory conditions. Indeed, genetic ablation or pharmacological inhibition of either only TRPV1 (10–13) or only TRPM3 (14–16) is sufficient to fully suppress inflammatory heat hyperalgesia. However, the precise mechanisms and the relative contributions of the heat-activated TRP channels to nociceptor sensitization under inflammatory conditions remains incompletely understood.

To address this problem, we induced experimental inflammation in one hindpaw of mice, and evaluated changes in expression of the three heat-sensitive TRP channels in sensory neurons. By combining retrograde labelling and quantitative *in situ* hybridization using RNAscope, we demonstrate a significant increase in mRNA encoding TRPM3 in dorsal root ganglion (DRG) neurons innervating the inflamed paw. Furthermore, we developed GCaMP3-based confocal imaging of DRG neurons *in situ*, which allowed us to measure inflammation-induced changes in the activity of the heat-sensitive TRP channels, both in the neuronal cell bodies and in the intact nerve endings in the skin. These experiments revealed that tissue inflammation provokes a pronounced increase in the activity of nociceptors co-expressing TRPM3 with TRPV1 and TRPA1 channels. Notably, pharmacological inhibition of TRPM3 not only eliminated TRPM3-mediated responses, but also normalized TRPV1- and TRPA1-mediated responses in neurons innervating the inflamed paw. These findings elucidate a central role of TRPM3 in inflammatory hyperalgesia, and provide a solid base for the development of TRPM3 antagonists to treat inflammatory pain.

## Results

### Increased expression of TRPM3 mRNA in sensory neurons innervating inflamed tissue

To address the role of heat-activated TRP channels in inflammatory heat hyperalgesia, we used a mouse model of complete Freund’s adjuvant-induced (CFA-induced) peripheral inflammation. In this established model, unilateral hind paw injection of CFA produces a strong inflammation associated with pronounced heat hyperalgesia, and the contralateral hind paw can be used as an internal control.

First, we addressed whether tissue inflammation is associated with increased expression of mRNA encoding heat-activated TRP channels, in particular in sensory neurons that innervate the inflamed tissue. Sensory neurons innervating the hind paws of mice have their cell bodies in the dorsal root ganglia, primarily at the lumbar levels L3-L6. These ganglia also contain cell bodies of sensory neurons that innervate other parts of the body, including viscera, necessitating an approach to specifically label neurons that have endings in the hind paw. Therefore, we intraplantarly injected the retrograde label WGA-AF647 in both hind paws seven days prior to tissue isolation, which resulted in an effective labelling of cell bodies of sensory neurons that innervate the injected territory (Figure 1A). We used single-molecule fluorescent RNA *in situ* hybridization (RNAscope) (17), to quantify the levels of mRNA encoding TRPM3, TRPA1 and TRPV1 in the ipsi- and contralateral DRGs (Figure 1B). We compared the levels of mRNA expression, both between retrogradely labelled (WGA-AF647^+^) and unlabelled (WGA-AF647^−^) neurons on the ipsilateral side, and between WGA-AF647^+^ neurons on the ipsi-versus contralateral side. Importantly, this analysis revealed a highly significant increase in the TRPM3 mRNA levels in the ipsilateral, WGA-AF647^+^ DRG neurons, both when compared to WGA-AF647^+^ neurons on the contralateral side, and to WGA-AF647^−^ neurons on the ipsilateral side (Figure 1C). These results provide strong evidence that inflammatory heat hyperalgesia is associated with increased transcription of the molecular heat sensor TRPM3, specifically in sensory neurons innervating the inflamed tissue. In the case of TRPV1, we did not detect a significant difference in mRNA levels between WGA-AF647^+^ neurons in the ipsilateral versus contralateral DRG neurons, arguing against a specific inflammation-induced expressional upregulation (Figure 1D). Finally, we did not observe significant inflammation-related changes in mRNA levels of TRPA1.

**Figure 1.**
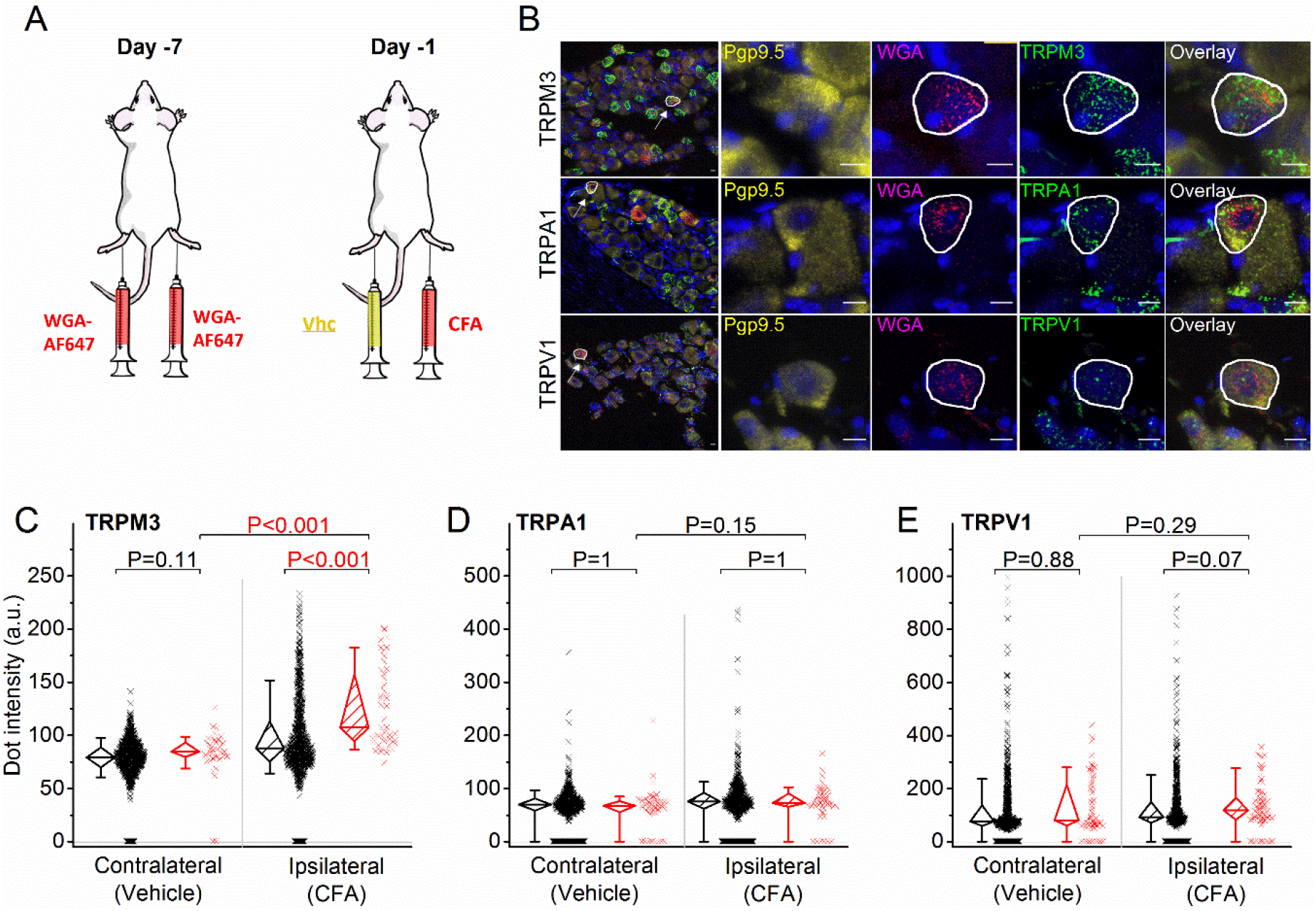
RNA expression of heat-activated TRP channels in sensory neurons innervating inflamed and control hind paws. **A,** experimental setting: seven days before analysis, mice were injected bilaterally with the retrograde label WGA-AF647. One day before imaging, the animals received a CFA injection in one paw, and vehicle in the contralateral paw. **B,** representative fluorescent images of processed L5 DRG. Shown are RNAscope stainings for the neuronal marker Pgp9.5 (yellow), along with specific probes for TRPM3 (top), TRPA1 (middle) or TRPV1 (bottom) (green). Retrogradely labelled neurons are identified based on the WGA-AF647 staining (magenta), and the blue color represents the nuclear marker DAPI. The leftmost and rightmost images represent overlays of the entire ganglion or the indicated cell, respectively. **C-E,** quantification of the total intensity of RNAscope dots per DRG neuron, comparing retrogradely labelled (red) and unlabelled (black) sensory neurons, from the contralateral and ipsilateral L3-L6 DRG. Statistical comparisons between groups were made using Kruskal-Wallis ANOVA with Dunn’s posthoc test. Data are from 6 mice. The total numbers of analysed neurons were for TRPM3: 752 ipsilateral and 1299 contralateral; for TRPA1: 954 ipsilateral and 947 contralateral; for TRPV1: 1054 ipsilateral and 995 contralateral.

### TRPM3 drives increased TRP channel responses in cell bodies of DRG neurons

Next, we investigated whether sensory neurons innervating inflamed tissue exhibit altered functionality of heat-sensitive TRP channels. In a first analysis, we focused on functional expression of TRP channels in the neuronal cell bodies. Since procedures to isolate and culture DRG neurons significanltly alter the expression levels of many ion channels (18), we developed an assay where the DRG was imaged as a whole *in situ,* using spinning-disk confocal imaging. For these experiments, we made use of a mouse line that expresses the genetically encoded calcium sensor GCaMP3 in TRPV1-lineage neurons (TRPV1-GCaMP3 mice), which include all the neurons involved in thermosensation and nociception (19, 20). Retrograde labelling using WGA-AF647 (see above) was used to identify the neurons that innervate the mouse paw, and changes in GCaMP3 fluoresence were monitored upon application of specific agonists for TRPV1 (capsaicin), TRPA1 (mustard oil; MO) and TRPM3 (the combination of pregnenolone sulfate; PS and CIM0216), and of a depolarizing high K^+^ solution resulting in TRP channel-independent Ca^2+^-influx via voltage-gated Ca^2+^ channels (Figure 2A). The efficiency of the retrograde labelling was similar in the ipsi- and contralateral side, with 25.8 ± 9.7 % and 27.7 ± 5.7 % of the neurons being WGA-AF647^+^, respectively. Both in the ipsi- and contralateral DRG, we observed calcium responses to the different agonist applications, which were indicative of neurons with different patterns of functional expression of one, two or all three heat-activated TRP channels (Figure 2B,C). We did not observe any significant differences in the response to different TRP channel agonists between WGA-AF647^+^ and WGA-AF647^−^ neurons in the contralateral DRG, indicating that the retrograde label by itself did not affect TRP channel activity at the level of the cell bodies (Figure 2D-F). Importantly, responses to TRPM3 agonists were substantially larger in the ipsilateral WGA-AF647^+^ neurons compared to both the ipsilateral WGA-AF647^−^ neurons and to the WGA-AF647^+^ contralateral neurons, indicating a specific functional upregulation of the channel in neurons innervating the inflamed paw (Figure 2D). In the case of TRPA1 and TRPV1, we also found a significant increase in responses in the ipsilateral WGA-AF647^+^ neurons compared to the WGA-AF647^+^ contralateral neurons (Figure 2E,F).

**Figure 2.**
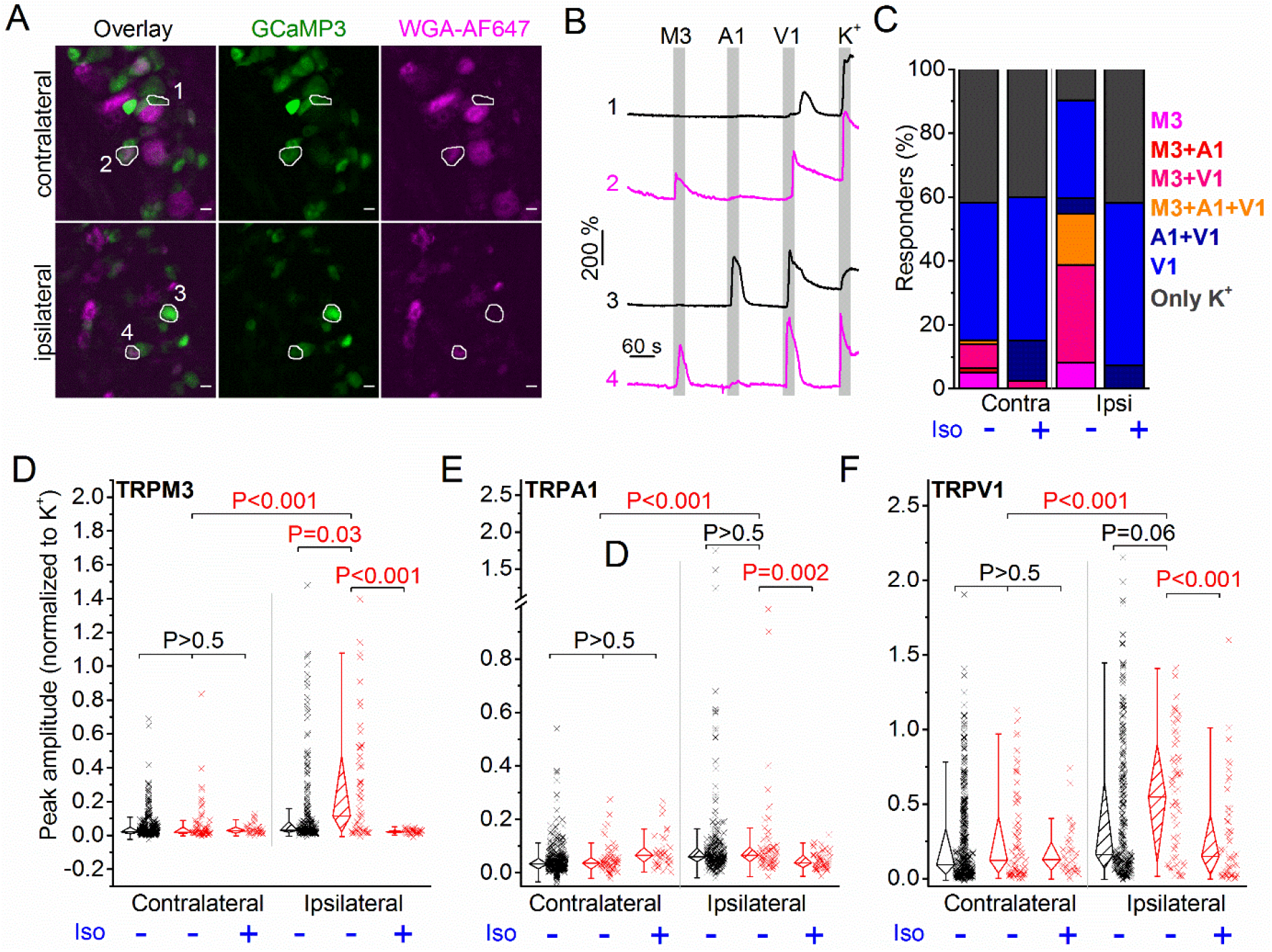
Inflammation-induced changes in TRP channel activity in DRG cell bodies. **A**, confocal images of the ipsi- and contralateral L5 DRG of a CFA-treated mouse showing GCaMP3 (green) and WGA-AF647 (magenta) fluorescence. **B,** representative examples, corresponding to cells indicated in panel (**A**) of changes in GCaMP3 fluorescence (ΔF/F_0_) in response to application of agonists of TRPM3 (PS + CIM0216; M3) TRPA1 (MO; A1) and TRPV1 (capsaicin; V1) and of a depolarizing high K^+^ solution (K^+^). **C,** percentage of neurons responding to the indicated (combinations of) agonists in retrogradely labelled DRG neurons from the ipsi- and contralateral sides, in the absence and presence of the TRPM3 antagonist isosakuranetin (iso). Cells that did not respond to high K^+^ stimulation were excluded from the analysis. **D-F,** peak amplitudes of responses to the TRPM3, TRPA1 and TRPA1 agonists, normalized to the response to the depolarizing high K^+^ solution, comparing retrogradely labelled (WGA-AF647^+^, red) and unlabelled (WGA-AF647^−^, black) neurons on the ipsi- and contralateral side. Where indicated, DRGs were incubated with isosakuranetin. Statistical comparisons between groups were made using Kruskal-Wallis ANOVA with Dunn’s posthoc test. Data are from 9 mice. Numbers of analysed neurons were 348 for ipsilateral and 526 for contralateral side.

However, there were no significant differences in the response amplitudes between WGA-AF647^+^ and WGA-AF647^−^ neurons on the ipsilateral side, suggesting that increased responsiveness to TRPA1 and TRPV1 agonists may not be limited to neurons innervating the hind paw. Notably, in the population of neurons that innervate the inflamed paw we observed an pronounced increase in the fraction that functionally co-expresses TRPM3 with TRPV1 and TRPA1. Further, the fraction of neurons excitable with TRPM3 agonist was also highly increased (Figure 2C).

Neurons innervating the inflamed paw showed a robust increase in mRNA expression of TRPM3, but not of TRPV1 and TRPA1 (Figure 1). We therefore considered the possibility that increased TRPM3 expression and activity provoked by inflammation may contribute to enhanced excitability of sensory neurons, resulting in enhanced responses to TRPA1 and TRPV1 agonists. To investigate this possibility, we tested the responses of WGA-AF647^+^ DRG neurons to TRP channel agonists in the presence of the TRPM3 antagonist isosakuranetin (21). As expected, isosakuranetin effectively eliminated the responses to TRPM3 agonists, illustrating the suitability of this compound to study TRPM3 functionality in DRG (Figure 2C,D). In WGA-AF647^+^ neurons on the contralateral side, isosakuranetin had no effect on the responses to MO or capsaicin (Figure 2E,F), in line with the reported selectivity of the antagonist for TRPM3 (21). Importantly, isosakuranetin caused a significant inhibition of the responses to capsaicin and MO in WGA-AF647^+^ neurons on the ipsilateral side, which were brought back to a level similar to the control (uninflamed) state (Figure 2E,F). Likewise, the fraction of neurons responding to TRPV1 and TRPA1 agonists in the ipsilateral, WGA-AF647^+^ neurons was restored to the control level in the presence of isosakuranetin (Figure 2C).

Taken together, these results indicate that inflammatory heat hyperalgesia is associated with increased functional expression of all three heat-activated TRP channels at the level of the DRG cell bodies. Notably, pharmacological inhibition of TRPM3 not only eliminates the responses to TRPM3 agonists, but also normalizes the responses to TRPA1 and TRPV1 agonists. These findings are in line with a model where increased molecular and functional expression of TRPM3 in the context of tissue inflammation enhances the excitability of sensory neurons, contributing to augmented responses to TRPA1 and TRPV1 agonists.

### Optical measurement of TRP channel activity in cutaneous nerve endings

Whereas these results indicate increased functionality of heat-activated TRP channels in the cell bodies of sensory neurons innervating the inflamed paw, they do not provide information regarding changes at the level of the sensory nerve endings. To address this issue, we developed an approach that allows direct measurement of TRP channel activity in intact cutaneous peripheral nerve endings in mouse hind paw skin (Figure 3A). In this assay, a skin flap of the hind paw and the innervating saphenous nerve (but lacking the DRG cell bodies) of TRPV1-GCaMP3 mice was excised and fixed in an organ bath, corium side up. Calcium signals in nerve endings expressing GCaMP3 were visualized from the epidermal side using a spinning-disk confocal microscope, while TRP channel agonists or a depolarizing high K^+^ solution were locally applied using a rapid perfusion system (Figure 3A). We adapted a published algorithm to automatically detect contiguous regions of interest (ROIs) exhibiting synchronous activity, which we interpret as individual branches of sensory nerve endings (22). As illustrated in Figure 3B,C, this approach revealed distinct calcium responses in localized regions, indicative of sensory endings that functionally express TRPM3, TRPA1 and TRPV1. We used this novel approach to compare TRP channel activity in the skin of CFA- and vehicle-treated paws (Figure 4A).

**Figure 3.**
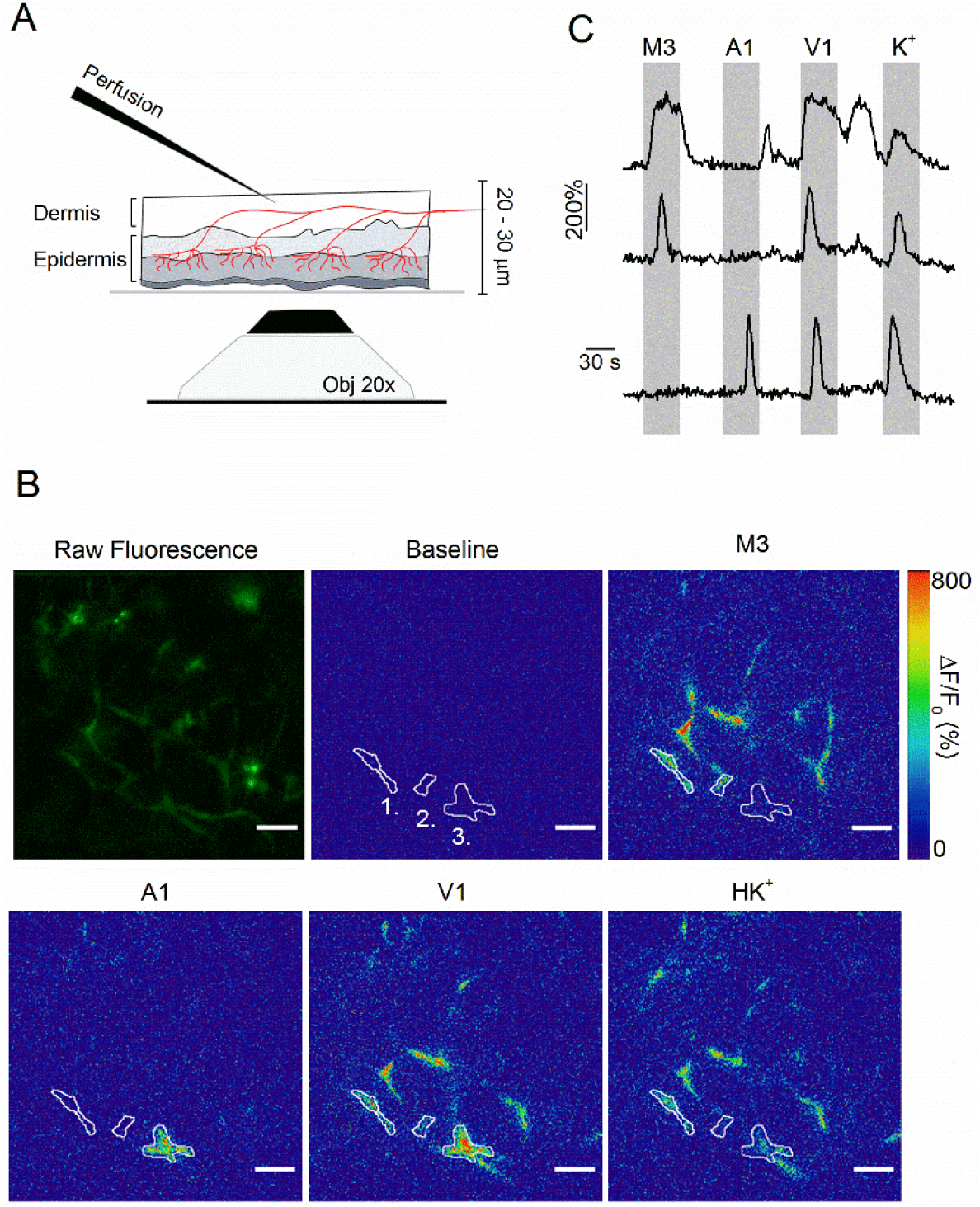
Optical measurement of TRP channel activity in peripheral sensory nerve endings. **A,** schematic illustration of the optical imaging setup. Sensory nerve fibers (red) innervating the dermal and epidermal skin layers are visualized using 488 nm laser light and an inverted spinning disk confocal microscope (20x objective). To avoid barrier effect of the epidermis, solutions (at 37 °C) were applied to the internal side of the sample from above. A total thickness of 20 – 30 μm was captured. **B,** the first image depicts the summed raw fluorescence of the entire imaging experiment. The next five images represent normalized fluorescence (ΔF/F_0_) at baseline (before the first stimulus), upon stimulation with TRPM3, TRPA1 and TRPV1 agonists, and with the depolarizing high K^+^ solutions. Three automatically detected ROIs, corresponding to the traces in panel **C**, are indicated. Scale bar is 50 μm. See **Supplementary Movie 1**. **C,** time course of normalized GCaMP3 fluorescence (F/F_0_) from three different ROIs depicted in panel **B**, with indication of the application periods of TRP channel agonists.

**Figure 4.**
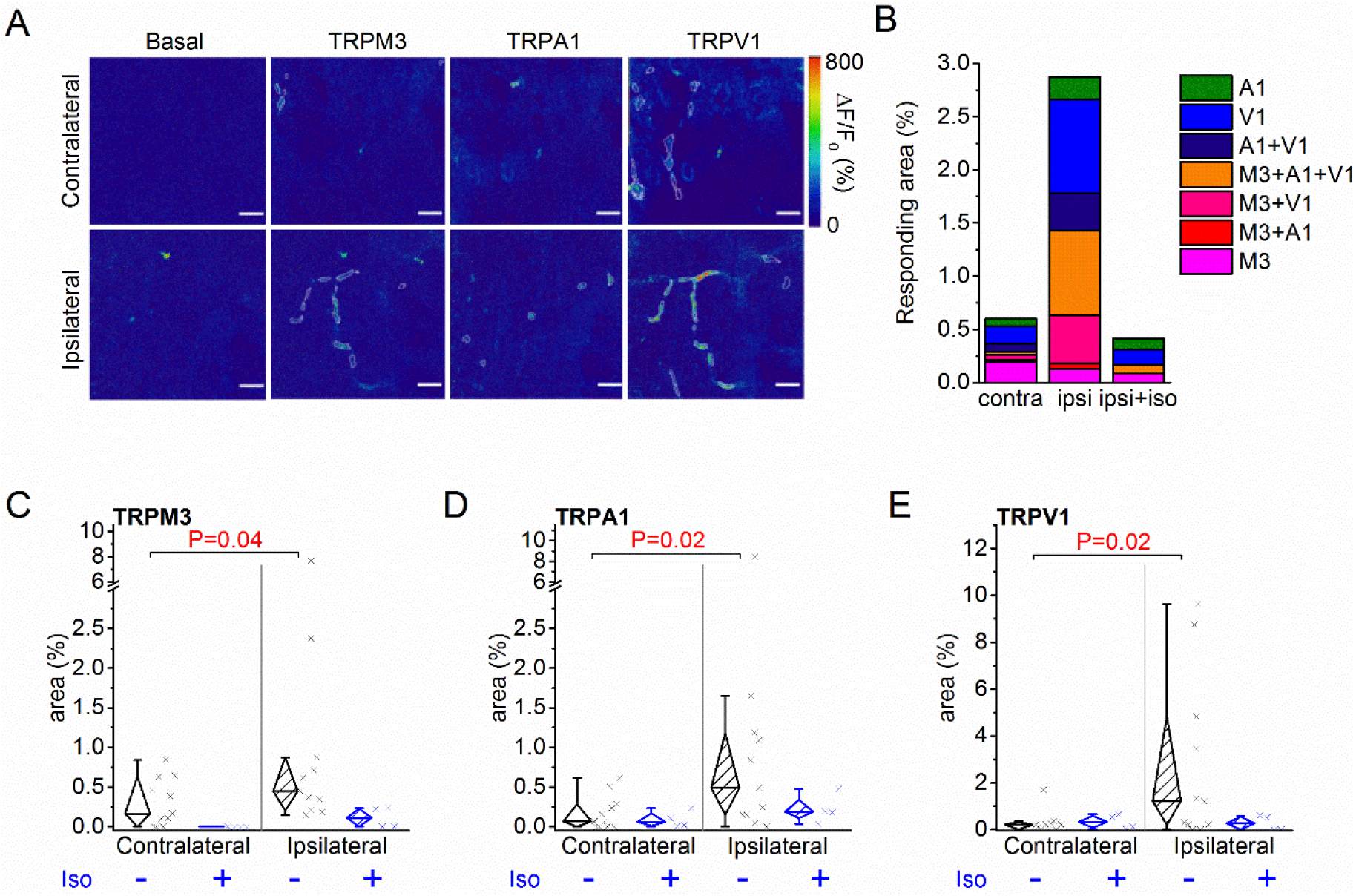
Increased TRPM3 activity in peripheral sensory nerve endings during inflammation. **A,** normalized fluorescence at baseline (before the first stimulus), and upon stimulation with TRPM3, TRPA1 and TRPV1 agonists. Scale bar is 50 μm. **B,** percentage of the total imaged area responding to the indicated (combinations of) agonists in the contra- and ipsilateral paws, and following isosakuranetin (iso) incubation. **C-E,** responsive areas to the indicated agonists in the contra- and ipsilateral skin, both in the absence and presence of isosakuranetin. The paired Wilcoxon Signed Rank Test was used for a paired comparison of the responsive area in the ipsi- and contralateral paw skin of 11 mice.

Since we measure responses from GCaMP3-positive nerve endings in the skin, not from clearly identifiable individual cells as in the DRG assay, a quantitative comparison of responses between ipsi- and contralateral paw posed several problems. First, the individual regions of interest that were automatically identified based on proximity and correlated activity represent only segments of individual sensory nerve endings, not the entire ending. Therefore, quantification of the percentage of nerve endings that respond to the specific TRP channel agonists was not feasible. Second, nerve endings in the skin did not always show robust responses to a depolarizing high K^+^ solution, even when they showed robust responses to one or more TRP channel agonists, making a normalization as was done for the neuronal cell bodies unreliable. The lack of consistent responses to a depolarizing high K^+^ solution may reflect that not all nerve endings contain voltage-gated Ca^2+^ channels or that inactivation of voltage-gated channels occurred due to the preceding TRP channel activation.

Therefore, to quantify the functional expression of the three TRP channels, we determined the total area of automatically detected ROIs that showed a correlated response to the specific agonists, normalized this active area to the total imaged area (Figure 4A), and made a paired comparison between the ipsi- and contralateral paws of the same animal. This analysis revealed increased reactivity to agonists of all three channels in the inflamed skin (Figure 4A-E), similar to the findings in the DRG cell bodies (Figure 2). In particular, we observed a pronounced increase in the surface of nerve endings exhibiting functional coexpression of TRPM3 with TRPV1 and TRPA1 (Figure 4B).

Following preincubation of CFA-treated skin with isosakuranetin, responses to TRPM3 agonists were largely abolished, whereas the surface of nerve endings responding to the TRPA1 and TRPV1 agonists returned to the level of the vehicle-treated paw (Figure 4B-E). These results represent, to our knowledge, the first direct observation of functional upregulation of heat-activated TRP channels in the context of inflammation in intact nerve endings in the skin, and indicate that the functionality of these channels is similarly affected in the nerve terminals and cell bodies.

## Discussion

In recent work, we demonstrated that three TRP channels (TRPM3, TRPV1 and TRPA1) act as redundant sensors of acute heat: neuronal heat responses and heat-induced pain are preserved in mice in which two of these three TRP channels were genetically ablated, but combined elimination of all three channels fully abolishes the withdrawal reflex from a noxious heat stimulus (9). Notably, earlier work had also revealed an absolute requirement of both TRPV1 and TRPM3 for the development of inflammatory heat hypersensitivity, since genetic ablation or pharmacological inhibition of either channel individually fully abrogates heat hyperalgesia in the rodent CFA model (10–16). These combined findings suggested that inflammation modulates the functional interplay between these heat-activated TRP channels leading to pathological heat hypersensitivity, but the underlying processes and mechanisms remained unclear. Here, by directly comparing the levels of mRNA expression and functional activity of TRPM3, TRPV1 and TRPA1 between sensory neurons innervating the healthy and inflamed hind paw in mice, we arrive at the following novel conclusions: (1) acute inflammation leads to a robust upregulation of TRPM3 expression at the mRNA level, specifically in DRG neurons innervating the inflamed paw; no significant changes were detected in the mRNA levels for TRPA1 or TRPV1; (2) inflammation is associated with increased functionality of TRPM3, TRPA1 and TRPV1 in DRG neurons, as evidenced by increased Ca^2+^ responses to specific agonists; this increased functionality is detected both in the neuronal cell bodies and in the nerve endings in the inflamed skin; (3) increased TRP channel activity in inflammatory conditions is associated with an increase in the fraction of cell bodies and nerve endings that functionally coexpress TRPM3 with TRPV1 and TRPA1, and (4) pharmacological inhibition of TRPM3 not only eliminates Ca^2+^ responses to TRPM3 agonists, but also normalizes responses to TRPV1 and TRPA1 agonists to the level of the non-inflamed paw. Taken together, these data indicate that inflammatory heat hyperalgesia is associated with a functional upregulation in the endings of nociceptors in the skin of all three molecular heat sensors implicated in the normal pain response to noxious heat, and point at TRPM3 as a key driver of hyperexcitability of nociceptors innervating inflamed tissue.

Whereas this is, to our knowledge, the first study demonstrating inflammation-induced alterations in TRPM3 expression at the mRNA level, earlier studies had already investigated alterations in TRPA1 and TRPV1 mRNA using the rodent CFA model. In line with our current results, several earlier studies report that transcript levels for TRPV1 were found unchanged after CFA treatment when assessed using quantitative PCR or RNA protection assays (23, 24). However, some earlier studies found increased levels of TRPA1 mRNA in lumbar DRG following CFA treatment of the hind paw (25, 26). These results seem at odds with our current findings, which did not reveal a statistically significant increase in TRPA1 mRNA in neurons innervating the inflammed paw. It should be noted, however, that in these earlier studies, bulk mRNA from entire ganglia was analysed, which inevitably includes not only neurons that innervate injured tissue, but also neurons from the same DRG that innervate healthy tissue, as well as non-neuronal cells present in DRG such as satellite glia and fibroblasts. In contrast, our approach using quantitative *in situ* hybridization and retrograde labelling allowed us to specifically measure mRNA levels in the cell bodies of the sensory neurons that innervate the inflamed or control hind paw. On the other hand, given the significant variability in TRPA1 mRNA levels between individual DRG neurons observed in our RNA-scope experiments, and the relatively limited number of neurons that were identified as WGA-AF647^+^, we consider the possibility that an overall increase in TRPA1 mRNA in the order of 30 % would not be revealed in our statistical analysis.

We used two novel approaches to assess inflammation-induced alterations in the functionality of the three TRP channels, one at the level of the cell bodies and the other at the level of the sensory nerve endings of DRG neurons. First, we developed an acute *ex vivo* preparation of the relevant part of the spinal column, where the cell bodies of sensory neurons were imaged within an intact DRG, but lacking the peripheral input. This preparation avoids the potential loss of specific neuronal populations as well as rapid alterations in their functional properties that are inherent to the isolation and culturing of DRG neurons (18). The use of WGA-AF647^−^labelling ensured the specificity for afferent neurons that innervated the hind paw tissue, whereas the use of the TRPV1-cre line to drive GCaMP3 expression allowed the specific recording from neurons involved in thermosensation and nociception. With the use of specific agonists, we were able to directly and quantitatively compare the TRPM3-, TRPA1- and TRPV1-induced activity between cell bodies of neurons that innervated the inflamed versus the control paw. Interestingly, we found increased responses to agonists for all three channels, and a particular increase in neurons that functionally co-expressed TRPM3 with TRPV1 and TRPA1. Increased activity of TRPV1 and TRPA1 in DRG neurons following CFA-induced inflammation is fully in line with earlier work (26–28), but this study is, to our knowledge, the first to reveal the specific increase in the subpopulation of DRG co-expressing the three heat-sensing TRP channels. Secondly, we developed an *ex vivo* assay, where we used GCaMP3-based calcium imaging to monitor TRP channel-mediated responses in intact sensory nerve endings in the paw skin. The skin preparation that we used is identical to the saphenous skin–nerve preparation that has been widely studied using extracellular electrodes to measure propagated action potentials from the receptive fields of single sensory nerve endings in the skin, including responses evoked by TRP channel agonists such as capsaicin (29, 30). This novel imaging approach allowed us to demonstrate for the first time that TRPM3, TRPA1 and TRPV1 show a pattern of partial spatial functional overlap in peripheral nerve endings in the skin. Moreover, similar to the findings in the neuronal cell bodies, we measured a pronounced increase in the surface of nerve endings that functionally express TRPM3 in the skin of the inflamed paw, particularly in endings that also respond to TRPV1 and TRPA1 agonists.

The potent TRPM3 antagonist isosakuranetin effecitively eliminated TRPM3-mediated calcium responses, both in the DRG cell bodies and nerve terminals. Intriguingly, isosakuranetin also tempered the responses to capsaicin and MO in neurons innervating the inflamed paw, restoring both the response amplitudes and the fraction of responding cells to the same level as in neurons innervating uninjured tissue. We can exclude the possibility that the reduced responses are due to a direct inhibitory effect of isosakuranetin on TRPV1 or TRPA1 activity, since the responses to capsaicin and MO were unaffected by the TRPM3 antagonists in neurons innervating healthy tissue. Likewise, earlier studies have shown that isosakuranetin does not inhibit heterologously expressed TRPV1 or TRPA1 (21). Instead, these data raise the possibility that increased molecular and functional expression of TRPM3 in neurons innervating inflamed tissue increases the excitability of nociceptors co-expressing TRPA1 and TRPV1, contributing to the augmented responses to agonist stimulation. This interpretation also provides a straightforward mechanism for the observation that heat hyperalgesia does not develop in TRPM3-deficient mice and is fully alleviated by TRPM3 antagonists (14, 16). Since heat hyperalgesia is also strongly attenuated by pharmacological inhibition or genetic ablation of TRPV1 (10–13), we hypothesize that those DRG neurons that gain functional coexpression of TRPM3 and TRPV1 under inflammatory conditions play a central role in the development of heat hypersensitivity.

In conclusion, the present findings provide the first evidence that TRPM3 expression and activity are increased in sensory neurons that innervate acutely inflamed tissue. In particular, we found a marked increase in sensory neurons that functionally coexpress TRPM3 with TRPV1 and TRPA1, two other TRP channels implicated in heat sensing and inflammatory pain, and this was observed both in the peripheral nerve endings and in the DRG cell bodies. Strikingly, pharmacological inhibition of TRPM3 reduced also normalized TRPV1- and TRPA1-mediated responses in nociceptors innervating the inflamed paw but not in the contralateral control neurons, suggesting that enhanced TRPM3 activity is an important driver of neuronal hyperexcitability under inflammatory conditions. Therefore, these results provide a straightforward rationale for the development of TRPM3 antagonists to prevent or alleviate inflammatory pain.

## Materials and Methods

### Animals

C57BL/6J wild type mice (Janvier Labs, Le Genest-Saint-Isle, France) and TRPV1-GCaMP3 mice on a C57BL/6J background were used. TRPV1-GCaMP3 mice were generated by crossing Rosa26-floxed-GCaMP3 mice (31) with TRPV1-cre mice (32). Mice were housed in a conventional facility at 21 ° C on a 12-hour light-dark cycle with unrestricted access to food and water. Both male and female mice between 8 and 12 weeks of age were used. Experiments were performed in concordance with EU and national legislation and approved by the KU Leuven ethical committee for Laboratory Animals under project number P075/2018 and P122/2018.

### Reagents

Reagents were purchased from Sigma-Aldrich (Chemical Co., St. Louis, Missouri) unless otherwise indicated.

### Retrograde labelling

To specifically label afferents from the hind paw skin, 10 μl Alexa Fluor 647-conjugated wheat germ agglutinin (WGA-AF647, Thermo Fisher Scientific, Invitrogen, Eugene, Oregon, USA; 0.8 % in sterile PBS) was injected intraplantar into both hind paws. The injections were performed 7 days prior to imaging. Initial experiments showed no edematous or hyperalgesic response to the retrograde label.

### Paw inflammation

Local inflammation was induced by injection of 10 μl complete Freund’s adjuvant (CFA, 1 mg/ml) into the plantar surface of the ipsilateral hindpaw of the studied mouse. The contralateral hind paw was injected with 10 μl vehicle (saline, Baxter, Lessen, Belgium). All ipsilateral mice hind paws showed substantial edema 24 hours after the injection, which was not observed in the control paw.

### Calcium imaging

Animals were euthanized using CO_2_ inhalation, and skin and DRG tissue were collected immediately.

#### Skin nerve preparation

Sensory nerve ending recordings were obtained from isolated dorsal hind paw skin preparations. The fur was removed with tape and the skin was gently dissected from the underlying tissue.

#### Isolation of DRGs

Bilateral L3-L6 DRGs were isolated. In brief, the spinal column was isolated, cleaned and split sagittally. The spinal cord, meninges covering the DRG, and the distal axon bundles were removed. Finally, a small segment of the spinal column containing the DRG of interest was extracted.

The isolated skin tissue and spinal column segments were maintained 1 hour on ice and 30 min at room temperature in synthetical interstitial fluid (SIF) solution, containing (mM): 125 NaCl, 26.2 NaHCO_3_, 1.67 NaH_2_PO_4_, 3.48 KCl, 0.69 MgSO_4_, 9.64 D-gluconic acid, 5.55 D-glucose, 7.6 Sucrose and 2 CaCl_2_. The pH was buffered to 7.4 using carbogen gas (95 % O_2_ and 5 % CO_2_). In some experiments, the latter 30 min incubation solution as well as the proceeding perfusion SIF solution contained TRPM3 blocker isosakuranetin (20 μM, Extrasynthese, Genay Cedex, France). During image acquisition, skin tissue was fixed with the corium side up in a glass-bottom microwell dish (MatTek, 35-mm petri dish, Ashland, USA). The spinal column sections were placed in the SIF-containing microwell dishes with the DRG facing towards the objective. Tissues were continuously superfused with 37 ° C SIF solution.

Calcium imaging was performed on an inverted spinning disk confocal microscope (Nikon Ti; Yokogawa CSU-X1 Spinning Disk Unit, Andor, Belfast, Northern Ireland), equipped with a 20x air objective (NA 0.8), a 488 nm laser light and a EMCCD camera (iXon3 DU-897-BV, Andor). For image acquisition and instrument control, Andor iQ software was used. A z-stack of 11 frames (total thickness of 20-30 μm) was captured consecutively during the entire measurement at a speed of 0.25 Hz. Agonists were diluted in SIF and applied using a heated perfusion system (Multi Channel Systems, Reutlingen, Germany). Before image acquisition, the DRG samples were excited at 640 nm to detect WGA-647^+^ retrogradely labelled cell bodies. The used stimuli to activate the specific TRP channels were for TRPM3: PS (100 μM) + CIM0216 (1 μM); for TRPA1: MO (100 μM); and for TRPV1 capsaicin (1 μM). Compounds were diluted in SIF supplemented with 0.1 % DMSO. A depolarizing high K^+^ solution in which all NaCl was replaced by KCl was applied at the end of each experiment to identify excitable cells. All solutions were applied at 37 ° C and administered to the immediate proximity of the recording field.

### Image processing and analysis

Recordings were first z-averaged in Fiji (imageJ, 1.52i) and corrected for translational drift using a reference image that averaged the first 10 frames prior to tissue stimulation (Turboreg plugin, imageJ).

#### Skin nerve

After drift correction, we used custom-made routines in Igor (Wavemetrics) to subtract fluorescence background. The background was obtained by fitting a fourth-order polynomial surface to non-responsive areas, which were defined as those pixels where the coefficient of variation in time was below a threshold, which was automatically obtained following the Otsu algorithm. After background correction, we used the ‘constrained non-negative matrix factorization for micro-endoscopic data’ framework (CNMF-E, Matlab R2017b) to identify ROIs as contiguous areas with temporally correlated activity (22). The background-subtracted fluorescence was normalized to the basal fluorescence of the reference image (first 10 frames prior to stimulation), yielding F/F_0_ values.

#### In situ DRG

These recordings were analyzed using a general analysis protocol (GA2) in NIS-elements (NIS 5.20.00, Nikon Instruments Europe B.V.). Retrogradely labelled (WGA-647+) and unlabelled (WGA-AF647^−^) neurons were identified manually. Raw fluorescent traces were converted to F/F_0_, where F is the fluorescence at a certain time point of interest and F_0_ is the baseline fluorescence before stimulation. Cells that did not respond to high K^+^ where excluded from all further analysis.

To determine responders in the skin-nerve and DRG recordings, two criteria had to be fulfilled: first, the peak F/F0 during stimulation had to exceed 5 times the standard deviation of the F/F0 value before stimulation. Second, the peak of the first derivative of the fluorescent signal (dF/dt) had to exceed the standard deviation of the dF/dt signal before stimulation.

### *In situ* hybridization

*In situ* hybridization was performed on 10-μm-thin cryosections of DRG neurons innervating the contralateral and ipsilateral hind paw. DRG tissue was isolated as described previously and immersed in 10 % neutral buffered formalin immediately after isolation. Prior to isolation, retrograde labelling and inflammation were induced as described previously. RNA transcripts were detected using the RNAscope 2.0 assay according to the manufacturer’s instructions (Advanced Cell Diagnostics, Hayward, CA, United States). Probes for mTrpv1 (cat number: 313331), mTrpm3 (cat number: 459911) and mTrpa1 (cat number: 400211) were purchased from Advanced Cell Diagnostics. The staining was performed using the RNAscope Fluorescent Multiplex Reagent Kit (cat number: 320850). As a positive control for neuronal cells, Pgp 9.5 staining was used. All cells were stained with DAPI and mounted on the slide with Gold Antifade Mountant. A total of 5802 neurons were analyzed from 12 lumbar (L5) DRG neurons (6 contralateral and 6 ipsilateral), isolated from 6 mice.

Slides were imaged using a Nikon NiE - Märzhäuser Slide Express 2 equipped with a Hamamatsu Orca Flash 4.0 in combination with a Plan Apo 40x (NA 0.95) and a custom made JOBS-GA2 protocol for sample detection. For analysis a GA3 script in NIS-Elements 5.20.00 was used. Cells were segmented manually. Dots in the green channel were detected using a rolling ball filter (1 μm) and spot detection (0.8 μm). From these dots the sum intensity and the number of dots per cell were measured. For the cells, the area, the XY coordinates and mean intensity in the green (RNA probe) and the far-red channel (WGA-AF647) were measured.

### Statistical analysis

Data analysis was performed using Origin software (OriginPro 2019b). Individual data points are displayed, along with box plots showing the median, first and third quartiles and outliers as whiskers. Since none of the relevant data were normally distributed, only non-parametric tests were performed. Specific tests are indicated in the legends. No statistical methods were used to predetermine the number of animals used in this study; since no a priory data were available for the novel imaging approaches used in this work, a power calculation was not feasible. However, our sample sizes are similar to those generally employed in other studies in the field, and potential limitations due to insufficient power are discussed in the text. Imaging and data analysis was performed without knowledge of the treatment (vehicle *versus* CFA).

## Acknowledgement

We acknowledge the Cell and Tissue imaging cluster (CIC; KU Leuven), where confocal microscopy was performed, and the Light Microscopy and Imaging Network (LiMoNe; CBD-VIB), where *in situ* hybridization slides were imaged. Calcium imaging at CIC was performed on an Andor Revolution Spinning Disk System supported by Hercules AKUL/09/50 to P.V.B. This research was further supported by grants from the VIB, KU Leuven Research Council (C1-TRPLe to T.V.), the Research Foundation-Flanders (FWO G0B7620N to T.V.) and the Queen Elisabeth Medical Foundation for Neurosciences (to T.V.).

## Notes

### Competing Interest Statement

The authors have declared no competing interest.

